# *naturaList*: a package to classify occurrence records in levels of confidence in species identification

**DOI:** 10.1101/2020.05.26.115220

**Authors:** Arthur Vinicius Rodrigues, Gabriel Nakamura, Leandro Duarte

## Abstract

1. There is a big volume of occurrence records available in biodiversity databases, but researchers should guarantee its quality before use it in scientific studies. A problem that might compromise the quality of occurrence data is species misidentification. We address this issue by presenting *naturaList*, a R package designed to classify species occurrence data according to identification reliability.
2. *naturaList* allows to classify species occurrences up to six levels of confidence in species identification, and to filter occurrence data accordingly. The highest level of confidence is assigned to records identified by a specialist, whose name must be provided by the user. The other five levels of confidence are derived from the occurrence data. We demonstrate *naturaList* functions using occurrences of *Alsophila setosa*, a tree fern species from Atlantic Forest, as example. We classified and filtered data in grid cells in order to maintain only the highest-level records in each cell. Then we selected only those records classified in the two highest levels of confidence.
3. From 323 occurrences of *Alsophila setosa* displaying geographic coordinates, 69 (21%) were identified by a specialist. After filtering the highest-level records inside grid cells, 102 records remained. From these grid cell filtered data, 38 occurrences (37%) were classified into the highest confidence level. Three records were removed using an interactive map module, due to falling in sea sites or outside the native range size of the species. Since we selected only records classified in the two highest levels of confidence, the final dataset contained 94 occurrence records.
4. *naturaList* guarantees the reproducibility of occurrence data processing and cleaning. Macroecologists, biogeographers and taxonomists might benefit from using *naturaList* package to evaluate the quality of species identification in occurrence data and by identify sites that need evaluation of taxonomic classification of species.

## Introduction

Open biodiversity databases (e.g. Global Biodiversity Information Facility – GBIF) provide data essential to studies of global biodiversity patterns. However, their use imposes big challenges regarding the quality of occurrence records data (Yesson et al., 2007; Anderson, 2012; Serra-Diaz, Enquist, Maitner, Merow, & Svenning, 2017). Solutions to problems related to georeferencing of occurrence records (Yesson et al., 2007) have gained attention and some reproductible solutions (Serra-Diaz et al., 2017; Zizka et al., 2019) have been presented. Undoubtedly, georeferencing quality is an important issue in species occurrence records; nonetheless, the reliability of species identification itself is paramount to guarantee data quality (Anderson, 2012; Araújo et al., 2019; Feng et al., 2019). This problem lacks automated processes to cope with, which is ideal to deal with big data. To assure for the quality of species identification remains being a subjective task, usually depending on expert taxonomists, or worst, being ignored.

There are some challenges in detect species identification errors from biodiversity databases. First, the user must have taxonomic skills to perform manual checking, and it is possible only if the occurrence records have images associated to them; nonetheless, even for experts, images are not always enough to confirm species identity. Second, depending on the amount of occurrence records, manual checking could be very time consuming, or even impossible to conclude. The presence of misidentification in datasets used in Species Distribution Models (SDM) may impact the performance and prediction of the models, enhancing the uncertainty in these SDM (Anderson, 2012; Araújo et al., 2019). Therefore, a method that account for such inaccuracies in the data by identifying and classifying them in levels of quality is of great importance because allows to cope with big data in a fast and reproductible way.

Thus, our aim is to present the *naturaList* R package, which provides tools for classifying occurrence records based on well-defined confidence levels of species identification. Further, *naturaList* enables to filter occurrence records within spatial grid cells based on the highest confidence level of species identification and provides an interactive module that allows to check out occurrence records in space and select them. We demonstrate package tools by applying *naturaList* functions to classify occurrence records of a tree fern species from South America.

## Materials and Methods

We propose a method for classification of the confidence in species determination based on the information provided by occurrence records databases. Our rationale is that the most reliable identification of a species is made by an expert taxonomist in the taxon (Araújo et al., 2019). To check if a species was determined by a specialist, *naturaList* user must provide a list of specialist names to be compared with the taxonomist name that determined the species. Moreover, other information is extracted from the occurrence dataset to produce subsequent confidence levels, detailed in the next section.

### *naturaList* R package

The *naturaList* package can be installed from github using devtools package (Wickham, Hester, & Chang 2019) with the code `devtools::install_github(“avrodrigues/naturaList”, built_vignette = TRUE`). The package has functions that allow the user to classify, filter and check species occurrences data, as described in the Table 1.

**Table 1.** Names and general description of the functions of the *naturaList* package.

| Function | Description |
| --- | --- |
|  | Classifies the occurrence records based on confidence in species |
| *classify_occ* | determination. |
| *create_spec_df* | Creates a specialist data frame from character vector. |
|  | Facilitates the search for non-taxonomist strings in the 'determined.by' |
| *get_det_names* | column of occurrence records dataset. |
|  | Filters the occurrence with most confidence in species identification |
| *grid_filter* | inside grid cells. |
| *map_module* | Checks out the occurrence records in an interactive map module. |

The classification in confidence levels of species identification is done with *classify_ occ* function. This function demands a data frame with the occurrence data and a data frame with specialist names. Each occurrence record is compared with the criteria from lower to higher levels of confidence and it is flagged with the highest criterion it met (Fig. 1). By default, *classify_occ* function uses six confidence levels (see Table 2), which, except for level 1, can be reordered according to the adequacy of the study. The output is a data frame containing the occurrence records with all the original information presented in the input dataset plus a column named ‘naturaList_levels’, indicating the confidence level of the occurrence record (codes are presented in Table 2).

**Figure 1.**
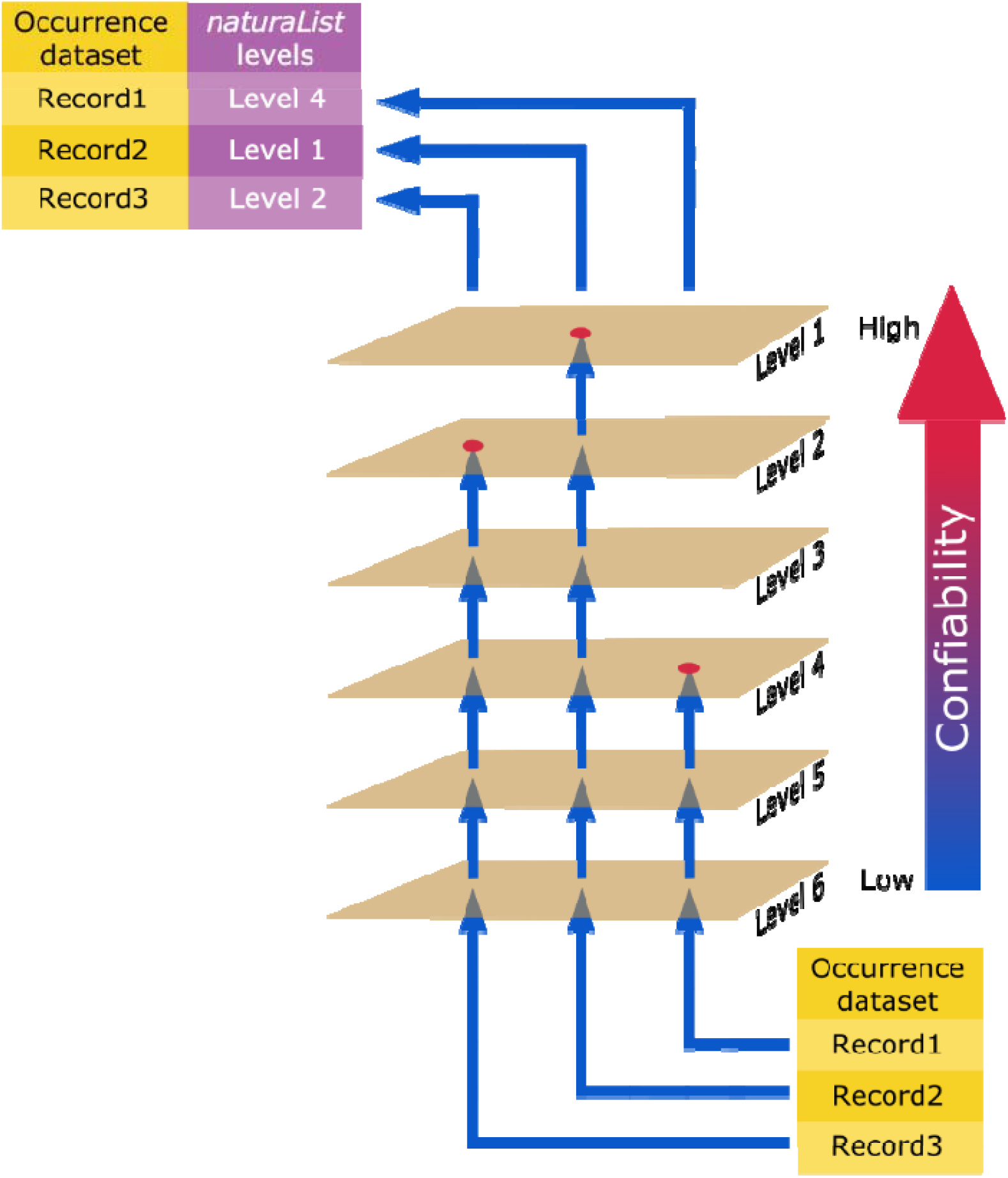
The scheme shows the classification procedure of occurrence records made by *classify_occ* function. Based on data frames of occurrence records and specialist names, *classify_occ* function checks for each occurrence record with the criteria of levels of confidence from lower to higher level and flags the occurrence record when the highest criterion it met.

**Table 2.** Levels of confidence in species determination used by *classify_occ* function. The level codes identify the criteria used and the numbers identify the order chosen by the user. The ordering presented in this table is the default order used in the function, which may be altered by the user. Level 1 is the highest level of confidence.

| Level code | Description | Default level<br>order |
| --- | --- | --- |
| det_by_spec | Species name is determined by a specialist in the<br>biological group, if not; | 1 |
| taxonomist | Taxonomist's name is provided, if not; | 2 |
| image | Occurrence has an associated image, if not; | 3 |
| sci_collection | Specimen is part of a scientific collection, if not; | 4 |
| field_obs | Occurrence is determined in a field observation,<br>if not; | 5 |
| no_criteria_met | No previous criteria are met. | 6 |

The occurrence dataset demanded by *classify_occ* must have at least the columns with the information showed in Table 3. By default, these column names follow the Darwin Core format (Wieczorek et al., 2012; https://dwc.tdwg.org/), thus data from databases that uses this format, e.g. GBIF, could be readily used in this function. Furthermore, other formats can be used by informing the corresponding column name in the function arguments. The specialist data frame demanded by *classify_occ* may be easily created from a character vector with *create_spec_df* function, which is ready to use in *classify_occ*.

**Table 3.**
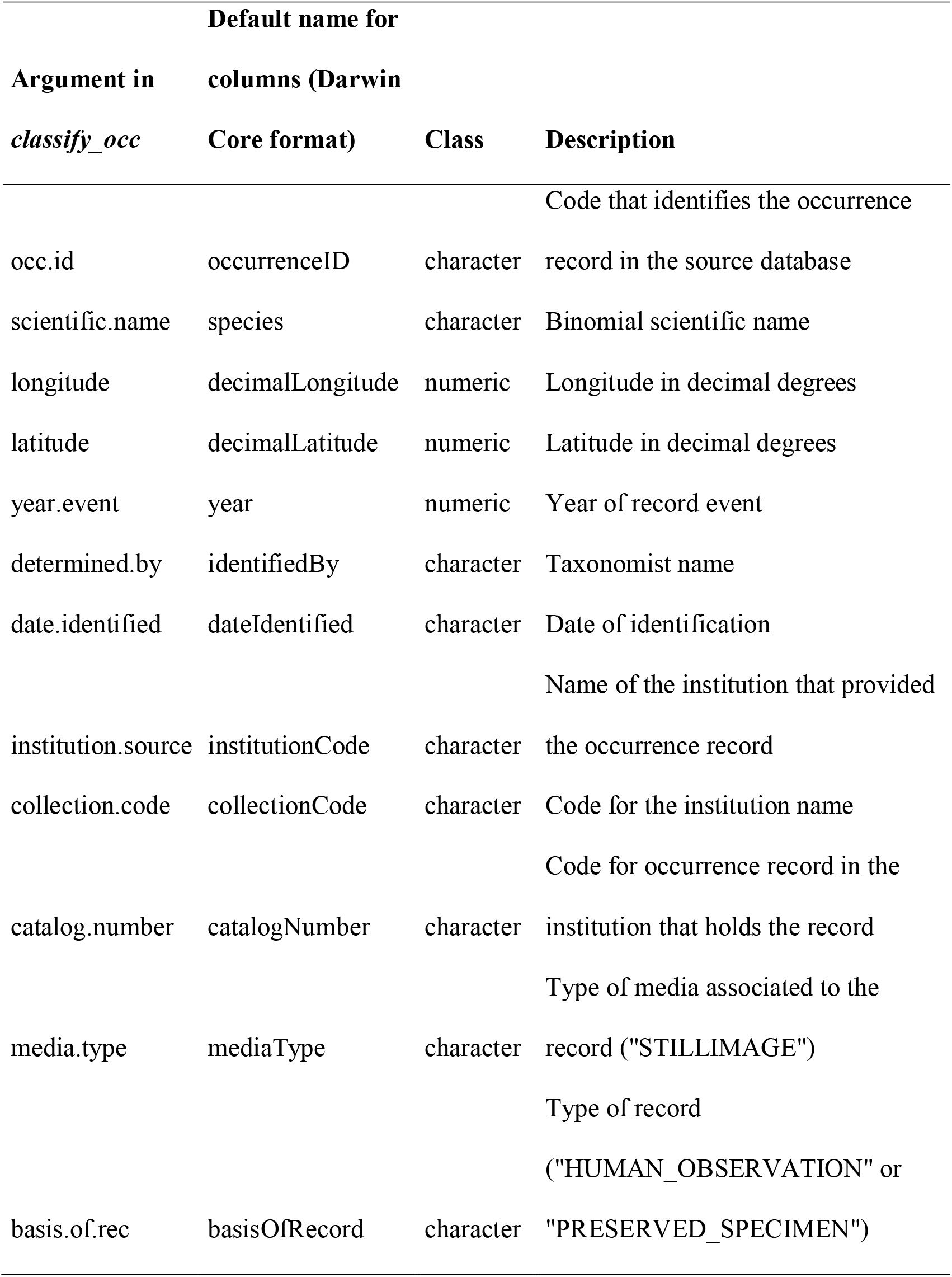
General format for occurrence records data frame used in *classify_occ* function.

To classify an occurrence record as identified by a specialist, the algorithm in *classify_occ* searches first if the surname of one or more specialist is present in the ‘determined.by’ column of the occurrence dataset. For those records that specialist surname matches to the names presented in ‘determined.by’ column the algorithm proceed in two ways: if the name in ‘determined.by’ column perfectly matches with specialist name, then the occurrence is classified at level 1. Otherwise, the function lists all strings that need verification by the user to assure it corresponds to a specialist name. In the manual verification, the user is asked by the function if the string printed in the R console corresponds (y), or do not (n) to a specialist’ name.

The classification process depends both on the quality of specialist dataset provided and the strings in the column indicated in the “determined.by’’ argument of *classify_occ* function. In some datasets, some strings may do not correspond to a valid name of a taxonomist (e.g. the string “unknown”). To overcome this problem, we suggest the use of *get_det_names* function to find strings that must be ignored as a taxonomist name. Based on this check out, the user should provide a character vector to *classify_occ* function in “ignore.det.names” argument, in order to ignore such strings that are not taxonomists’ names.

The *grid_filter* function selects the occurrence with highest level of confidence inside a spatial extent, given by a cell size defined by the user or from a raster layer. Thus, this function returns a data frame only with the best classified occurrence record of each cell. If two or more occurrences have the same confidence level in a cell, the function gives priority for occurrence that was 1) more recently determined; 2) more recently observed. If the equality persists or if a date was not provided, one of the occurrences is randomly chosen.

The *map_module* function is an interactive application that allows the user to manually filter records in the map and easily access the information of each record. This function could operate with output from *classify_occ* or *grid_filter* functions. In this map, the occurrence points are coloured according to the confidence levels classification. By clicking over a point, the user can see some information about that occurrence. When setting on the option ‘delete points with click’, clicking over a point works to delete that occurrence record. Furthermore, the user may select the classification levels to filter points or yet may draw a polygon to select occurrence in a region. These functionalities are helpful to find for spatial outliers and/or occurrences out of the native range (if the native range is known).

### Example

To demonstrate the use of *naturaList* package, we conducted a classification of the occurrence records of *Alsophila setosa*, a tree fern species from Atlantic Forest. For this, we downloaded all the occurrence records of *Alsophila setosa* species from GBIF (GBIF, 2019). We assumed that occurrence records had no issues regarding geographical coordinates and did not conduct any procedure to remove records before performing the classification procedure. To produce a list of specialists, we took the names of the authors of a checklist of ferns and lycophytes species of Brazil (Prado et al., 2015). Both datasets are available in the *naturaList* package.

We first conducted the classification of occurrences according to the six confidence levels, using default settings in *classify_occ*. Then we filtered the occurrence records in a grid cell with size of 0.5° with *grid_filter* function. The size of the grid cell was arbitrarily chosen only to show how to use the function. Finally, we checked if the occurrence points were inside the native range using *map_module* function. We considered information available in Flora do Brasil (http://floradobrasil.jbrj.gov.br), which inform the Brazilian states where species occur, and Plants of The World Online (http://www.plantsoftheworldonline.org/), which provides political regions where the species occur in South America, to define the native distribution range of *A. setosa*. The code used to produce this example is provided in the supporting information. Additionally, an introduction to the package can be found with the code: `vignette(“naturaList_vignette”)`.

## Results

The dataset of *Alsophila setosa* has 508 occurrence records. After using *classify_occ* the number of records decreased to 323 due to records without coordinates in the raw GBIF dataset, which were automatically removed by the default setting of *classify_occ*. The *classify_occ* function asked for manual checking of four strings for specialist names, which were further confirmed as true ones. From the 323 occurrences returned, 69 occurrences were classified as level 1, 218 as level 2, 12 as level 3 and 24 as level 4, which represented 21.4%, 67.5%, 3.7% and 7.4% of the total classified occurrence records, respectively.

The dataset with 323 classified occurrence records was then filtered in grid cells with 0.5° of size. This filtering reduced the dataset to 102 occurrences, in which 38 (37.3%), 58 (56.9%), 3 (2.9%), 3 (2.9%) were classified, respectively, in levels 1 to 4. The grid cell filtering enhanced the representativeness of occurrences classified in the level 1 (from 21.1 to 37.3%), by removing occurrences classified in lower levels that are placed in the same grid cell.

Finally, we used this dataset with 102 occurrences in the *map_module* to visually check for potential errors and to select only occurrence records in levels 1 and 2 (Fig. 2). First, we deleted three occurrence records, by setting on the button ‘delete points with click’ (Fig. 2b-II) and clicking on them; two of them were deleted because were placed in the sea and one because was out of the native range recognized by Flora do Brasil (the ‘x’ in Fig. 2b). Then, we selected the levels 1 and 2 to be maintained in the output dataset (Fig. 2b-I). The current selected records may be visualized above the ‘Done!’ button (Fig. 2b-III). Note that selection made in the Fig. 2b-IV only serves to display the points in the map. Polygons should also serve as spatial selection that the user can draw with tools in the left side of the map (Fig 2b-V). Finally, we clicked in ‘Done!’ button to assign the selected points to an R object. At the end of these procedure 94 occurrence records were maintained in the occurrence dataset.

**Figure 2.**
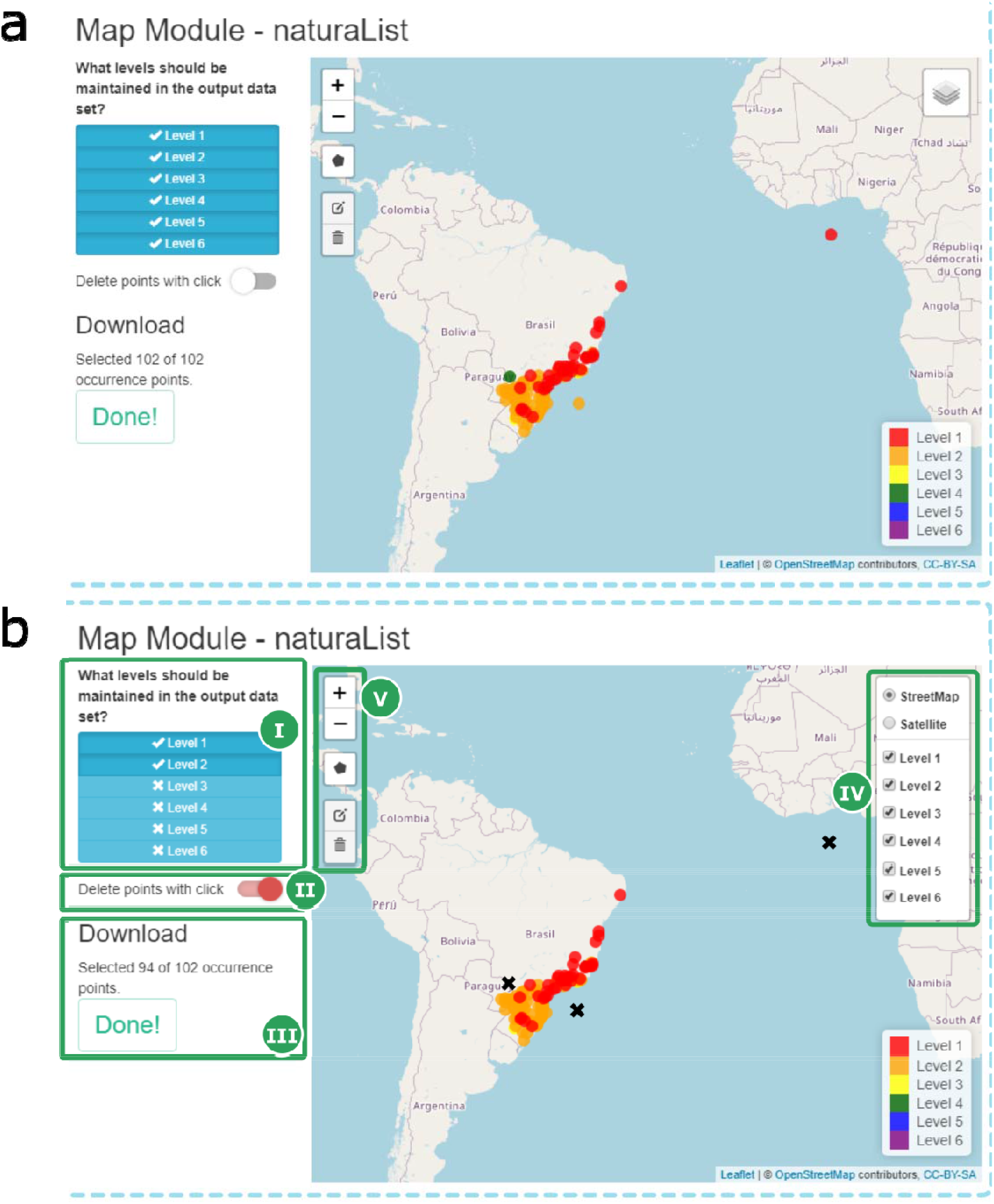
Features of *map_module* function used in the example with *Alsophila setosa* occurrence data. a) the initial display of occurrence records data; b) highlights the features of *map_module*. I – Select levels to be maintained in the output dataset; II – enable to delete point with click over occurrences; III – show how many points have been selected from the initial dataset; IV – display in the map only occurrences that belong to the selected levels, and turn visualization of the map between OpenStreetMap and Satellite; V – options to draw and edit polygons to select occurrence records that fall within polygon; x – highlights the deleted occurrence points.

## Discussion

Our example demonstrated that *naturaList* enables researchers to classify and quantify, based on well-defined criteria, the reliability of species identification in occurrence records from biodiversity databases. This may assist researchers to decide which levels of confidence is suitable to their study objective. Also, our example showed that the proportion of records with high quality can be improved after the use of *grid_filter* function, therefore it might enhance data quality to be used in SDMs. This increase, however, is dependent on the size of grid cell used in the filtering and the spatial distribution of records identified by specialists. The usefulness of such grid cell filtering is to evaluate the reliability of identification in the same scale of the environmental layer used to fit SDM. To this end, in the *grid_filter* the user may provide a raster layer, from which the function will use the cells to conduct the filtering.

Although SDM is the most popular application aiming to produce accurate estimates of species distribution, occupancy models have been also used for this purpose (Kéry, Gardner, & Monnerat, 2010). The classification of occurrence records regarding confidence in the identification might be useful for occupancy models that incorporate false positive probabilities (Royle & Link, 2006; Miller et al., 2011).

The tools described here could be also useful for taxonomists, that might identify areas in which there are occurrence records for a given taxon, but with uncertain reliability. Based on that information, the specialist could confirm (or not) the occurrence of the species – by visiting the areas or a scientific institution with preserved material – and thereby improving the quality of identification.

On one hand, the tools in *naturaList* enable researchers to account for the quality of species identification in big datasets as well as to report the steps used in the cleaning of data, which enhance the reproducibility of such studies. On the other hand, those tools might enhance the quality of species identification by guiding specialists in taxonomy to revise specimens with low determination confidence.

## Acknowledgments

We are thankful to the 2019 GBIF Ebbe Nielsen Challenge by recognize the importance of the tools described here. AVR is thankful to PhD grant from Coordenação de Aperfeiçoamento de Pessoal de Nível Superior - Brasil (CAPES) - Finance Code 001. LD research activities have been supported by CNPq Productivity Fellowship (grant 307527/2018-2). LD and GN research have been conducted in the context of the National Institutes for Science and Technology (INCT) in Ecology, Evolution and Biodiversity Conservation, supported by MCTIC/CNPq (proc. 465610/2014-5) and FAPEG (proc. 201810267000023).

## Author contributions

A.V.R. conceived the idea. A.V.R and G.N implement the package. All authors contributed in discussions, suggestions and in the writing of the manuscript.

## Data availability

The data used in the example and the package are available at https://github.com/avrodrigues/naturaList

