## Supplementary material for "*naturaList*: a package to classify occurrence records in levels of confidence in species identification": example_script.R

```
library(devtools)
install_github("avrodrigues/naturaList", built_vignette = T, force = TRUE)
library(naturaList)

data("A.setosa")
data("speciaLists")

# classification
occ.class <- classify_occ(A.setosa, speciaLists)
y
y
y
y
occ.class

# number of records by level
class.levels <- table(occ.class$naturaList_levels)
class.levels

# percentage of records by level
class.levels/sum(class.levels)*100

# filtering in grid cells
occ.grid <- grid_filter(occ.class)
occ.grid

# number of records by level
grid.levels <- table(occ.grid$naturaList_levels)
grid.levels

# percentage of records by level
grid.levels/sum(grid.levels)*100


# check out the occurence in the interactive map module
occ.selected <- map_module(occ.grid)
occ.selected

# number of records by level
selected.levels <- table(occ.selected$naturaList_levels)
selected.levels

# percentage of records by level
selected.levels/sum(selected.levels)*100
```
